# The Metabolomics of a Cysteinyl Leukotriene Receptor 1 (CysLTR1) Knock Out Mouse Model by Analysis of Bronchoalveolar Lavage Fluid Using Gas Chromatography-time of Flight Mass Spectrometry

**DOI:** 10.1101/2025.08.15.670145

**Authors:** Wilson Bamise Adeosun, Sibongiseni KL. Poswayo, Suraj P. Parihar, Du Toit Loots

**Affiliations:** Biomedical and Molecular Metabolism (BioMMet), North-West University, Hoffman Street, 2531, Potchefstroom, South Africa; Division of Medical Microbiology, Institute of Infectious Diseases and Molecular Medicine (IDM), Faculty of Health Sciences, University of Cape Town, Cape Town, South Africa; Centre for Infectious Disease Research in Africa (CIDRI-Africa), Faculty of Health Sciences, University of Cape Town, Cape Town, South Africa

**Keywords:** cysteinyl leukotriene receptor 1, bronchoalveolar lavage fluid, metabolomics

## Abstract

Cysteinyl leukotriene receptor 1 (CysLTR1), a potent lipid mediator, is known for its critical role in regulating inflammatory responses, particularly in asthma and airway diseases. While its function in immune cell recruitment has been previously reported, its broader impact on pulmonary metabolism remains largely unexplored. In this study, we investigated the metabolic consequences of CysLTR1 deletion in mice using GC-TOFMS-based metabolomics analysis of bronchoalveolar lavage fluid (BALF) from both CysLTR1 knockout (KO) and wild-type (WT) mice. The BALF from CysLTR1 KO mice exhibited significantly reduced levels of glucose, gluconic acid, sedoheptulose, D-xylose, glucosamine, glyceric acid, and 1-methylinosine, indicating impaired glucose uptake and dysregulation of glycolysis and gluconeogenesis. Further disruption of glucose-associated pathways, including the pentose phosphate pathway and purine metabolism, alongside reduced 1-methylinosine levels, suggests altered RNA turnover. In addition, decreases in butanoic acid, decan-2-ol, and 1-hexadecanol point to dysregulated fatty acid metabolism, potentially as a compensatory response to glucose deficiency. Altered levels of mandelic acid, glutaric acid, tricarballylic acid, and decan-2-ol, some of which are derived from corn-based diets also indicate changes in the pulmonary microbiome. Overall, the deletion of CysLTR1 significantly disrupts pulmonary metabolic homeostasis, affecting the metabolism of carbohydrates, lipids, amino acids, nucleotides, and microbial-derived metabolites.

## Introduction

Lipid mediators that are derived from the metabolism of the arachidonic acid through the 5-lipoxygenase (5-LOX) pathway are known as regulators of inflammation, vascular permeability, and smooth muscle contraction (1). Leukotrienes (LTs) and cysteinyl leukotrienes (CysLTs; LTC4, LTD4 and LTE4), are some of these lipid mediators that are synthesized through the 5-LO pathway. Leukotrienes, especially leukotriene B4 (LTB4), regulate how macrophages respond to antigens, resulting in increased inflammation. This is important in the progression of metabolic diseases including atherosclerosis (2), obesity and type 2 diabetes (3,4). With reference to the latter, the engagement of LTB4 to its receptor results in various inflammatory responses, leading to insulin resistance, and ultimately obesity and type 2 diabetes. Interestingly, elevated concentrations of LTB4 in type 2 diabetes patients, have also been shown to correlate with cardiovascular autonomic dysfunction (5).

Cysteinyl leukotrienes perform their biological functions through G-protein-coupled receptors, CysLTR1, CysLTR2, and CysLTR3. CysLTR1 has been intensively studied in allergic and asthmatic responses, but there is a growing body of evidence that this receptor has a broader role in regulating chronic metabolic dysfunction, because of its role in inflammatory regulation. For instance, CysLTs have been shown to increase the secretion of proinflammatory cytokines and adipokines, such as interleukin 6 (IL-6), monocyte chemotactic protein 1, tumor necrosis factor-α (TNF-α), nuclear factor kappa B (NF-κB), and macrophage inflammatory protein 1 (4,6). Therefore, targeting CysLTR1 and its ligands could be helpful in the management of inflammatory conditions besides asthma and allergic reactions.

Aside from classic metabolic diseases, CysTLR1 mediates inflammatory dysregulation during infections, such as in chronic infection with *Schistosoma mansoni*. It has been demonstrated that mice lacking CysLTR1 have reduced granuloma sizes, hepatic fibrosis, and liver enzyme release. This was also associated with increased anti-inflammatory cytokines. The authors also demonstrated that mice treated with montelukast (a CysLTR1 antagonist) in combination with praziquantel result in reduced cellular infiltration to the liver and reduced egg burden during chronic infection (7). Together, this data demonstrates that combinational therapy (CysLTR1 inhibition and praziquantel) can be used as a prophylactic to treat chronic schistosomiasis infection. This further highlights the broader role of CysLTs and their receptor (CysLTR1) as crucial mediators of inflammation and metabolic pathways in both infectious and non-infectious diseases, suggesting that CysLTR1 antagonists have the potential to be repurposed for resolution of metabolic inflammation and tissue destruction in various disease conditions and infections.

Bronchoalveolar lavage fluid (BALF) is an important biofluid used in the investigation of the lung’s microenvironment. The procedure involves washing of the lower respiratory tract (bronchoalveolar space) with saline solution which is then recovered. The solute content is composed mostly of alveolar macrophages, lymphocytes, neutrophiles, proteins and primary metabolites (8,9). A metabolomics analysis of BALF would subsequently provide a detailed snapshot of metabolic alterations that may have occurred in the pathophysiology of the lungs due to a perturbation. The many biological processes mice share with humans make them ideal models for biomedical research.

A rather recent but now a staple technology in biomedical research - metabolomics, comprehensively analyses metabolites associated with metabolic changes within a biological system at a given time. Gas chromatography-time of flight mass spectrometry (GC-TOFMS) is a widely used hybrid technique in metabolomics that combines the separation proficiency of GC with the detection capacity of MS, resulting in higher efficiency of metabolite identification in a sample.

This study investigated the role of CysLTR1 in the lung, by comparing the metabolic changes in a CysLTR1 KO mice model to a WT control, using GC-TOFMS metabolomic analysis of BALF samples. The results of this study expand our knowledge by providing additional valuable insights into the novel role of CysLTR1 in the lower respiratory system.

## Materials and Methods

### Sample collection

CysLTR1-deficient mice (Cysltr1-/-) were created by breeding heterozygous (Cysltr1-/+) animals on C57BL/6 background. Dr. Frank Austen from Harvard Medical School generously provided CysLTR1 heterozygous mice. These mice were housed in ventilated cages under specific-pathogen-free conditions at the research animal facility of the UCT Faculty of Health Science. Mice aged 8-12 weeks were used, with sex matching unless otherwise specified. The mice were terminally anaesthetized, after which a catheter was inserted into the trachea and secured in place. A 1 mL syringe containing sterile saline solution was then connected to the catheter and gently injected. The solution was gently aspirated while massaging the thorax of the mouse to collect bronchoalveolar lavage fluid (BALF). Approximately 500–800 µL of BALF per mouse was transferred into 2 mL cryotubes, immediately snap-frozen in liquid nitrogen, and stored at −80 °C until analysis. All animal studies adhered to the rigorous guidelines outlined in the South African National Standard for the Care and Use of Animals for Scientific Purposes (SANS 10386:2008). The study protocol was approved by the Animal Research Ethics Committee (AREC 022/024) at the Faculty of Health Science, University of Cape Town.

### Reagents

Deionised water from a Millipore MilliQ purification system was used throughout the study. Optima-grade acetonitrile was obtained from Fisher Scientific (Pittsburgh, USA). Unless otherwise stated, all other reagents and organic solvents used in this investigation were sourced from Sigma-Aldrich (St. Louis, MO, USA).

### Sample preparation and extraction

After adding 50 µL of the 3-phenylbutyric acid (Sigma-Aldrich) internal standard, the BALF samples were protein precipitated by adding 50 μL of acetonitrile, followed by centrifugation at 3000 rpm for 10 minutes at room temperature. A volume of 100 μL of each supernatant was subsequently evaporated to dryness before derivatization. The dried metabolic extract was oximated with 50 μL of methoxyamine hydrochloride in pyridine (15 mg/mL) (Merck, Darmstadt, Germany) at 50 °C for 1 h. Following oximation, 50 μL of N, O-bis (trimethylsilyl) trifluoroacetamide (BSTFA) with 1% trimethyl chlorosilane (TMCS) (Sigma-Aldrich, St.Louis, MO, USA) was added, and the mixture then incubated at 60 °C for 1 h to form trimethylsilyl (TMS) derivatives. The derivatized sample extracts were transferred to 2 mL glass vials containing 250 uL glass inserts.

One microliter of each sample extract was injected (1:1 split ratio) into a Pegasus BT GC-TOFMS instrument (Leco Corporation, St. Joseph, MI, USA), consisting of an Agilent 7890A gas chromatograph (Agilent, Atlanta, GA, USA) coupled to a time-of-flight mass spectrometer (TOFMS) (Leco Corporation, St. Joseph, MI, USA). Separation was achieved using Rxi-5-MS column (29.690 m, 0.25 mm internal diameter and 0.25 μm film thickness) (Restch GmbH & Co. KG, Haan, Germany). The front inlet temperature was held at a constant 270 °C, the transfer line temperature at a constant 250 °C, and the ion source temperature at a constant 200 °C, for the entire run. The initial GC oven temperature was set at 70 °C for 1 min followed by an increase in the oven temperature of 5 °C /min to a final temperature of 320 °C, at which it was held for 3 min. The detector acquisition delay for each run was 420 s, and offset with a filament bias of −70 eV. Spectra were collected from between 50 to 950 m/z at an acquisition rate of 20 spectra-per second. Mass spectral deconvolution, peak alignment and peak identification were performed using Leco Corporation’s ChromaTOF software (version 4.71). Mass spectral deconvolution, peak deconvolution, peak alignment and identification, were performed at a signal-to-noise ratio of 30, with a minimum of three apexing peaks. To eliminate the effect of retention time shifts and to create a data matrix containing the relative abundance of all compounds present in all samples, peaks with similar mass spectra and retention times were aligned using statistical compare, a package of ChromaTOF. Mass fragmentation patterns and their respective retention times were screened against commercially available (National Institute of Standards and Technology (NIST) 2020 and Whiley v12) and in-house libraries compiled from the mass spectra obtained from previously injected standards, for peak annotation, with a similarity setting of at least 70%. All metabolites markers identified as described in 2.5, where manually checked again by the analyst, by comparing their retention times and mass fragment patterns to that of the libraries, to confirm their identities before biological interpretation.

### Statistical analysis and metabolite identification

The relative concentrations of 201 compounds comprising the combination of quality control (QT), WT and KO samples were statistically analysed. Standard metabolomics data clean-up procedures were followed (including data pre-treatment, spectra deconvolution, missing data input, data normalization and batch effect correction), and the cleaned data was further processed using various univariate and multivariate statistical analyses (10,11). Multivariate models and univariate statistics were used complementarily, since univariate analysis examines each metabolite individually to identify significant changes while multivariate analysis takes into consideration the relationships between multiple metabolites simultaneously. Univariate statistics carried out included t-tests, fold change and effects size analyses, and carefully completed with the chemometric analysis of group differences using orthogonal partial least squares-discriminant analysis (OPLSDA). A principal component analysis (PCA) was also conducted to determine the compounds contributing most to the observed variability and their relationship to the groups. All data analyses were performed using Metaboanalyst 6.0 (12).

## Results

### Raw data summary

A total of 936 compounds were detected by the benchtop GC-TOFMS. The dataset was meticulously cleaned to eliminate “unknowns”, other artifact compounds of no biological relevance, and compounds without matching mass spectral entries in the libraries. This was followed by a merger of putative compounds with duplicate peaks such as monosaccharides and amino acids resulting in the aforementioned 201 compounds (excluding the internal standard) which were subjected to the aforementioned statistical analyses.

### Quality control

As indicated in Figure 1, clear grouping of the QC replicates indicates good instrumental stability and minimal batch effects throughout the analytical run. The consistent grouping of the QC samples supports the reliability of the data and suggests that machine drift or analytical variability is negligible. The QC samples, generated by pooling aliquots of the study samples encompass the overall metabolic variance across the study. All the BALF samples and QCs were randomized across and within batches prior to analysis, ensuring that any residual analytical variation was equally distributed among the groups, thereby minimizing potential bias. These observations affirm the quality and reproducibility of the dataset, validating its suitability for downstream multivariate and univariate statistical analyses.

**Figure 1.**
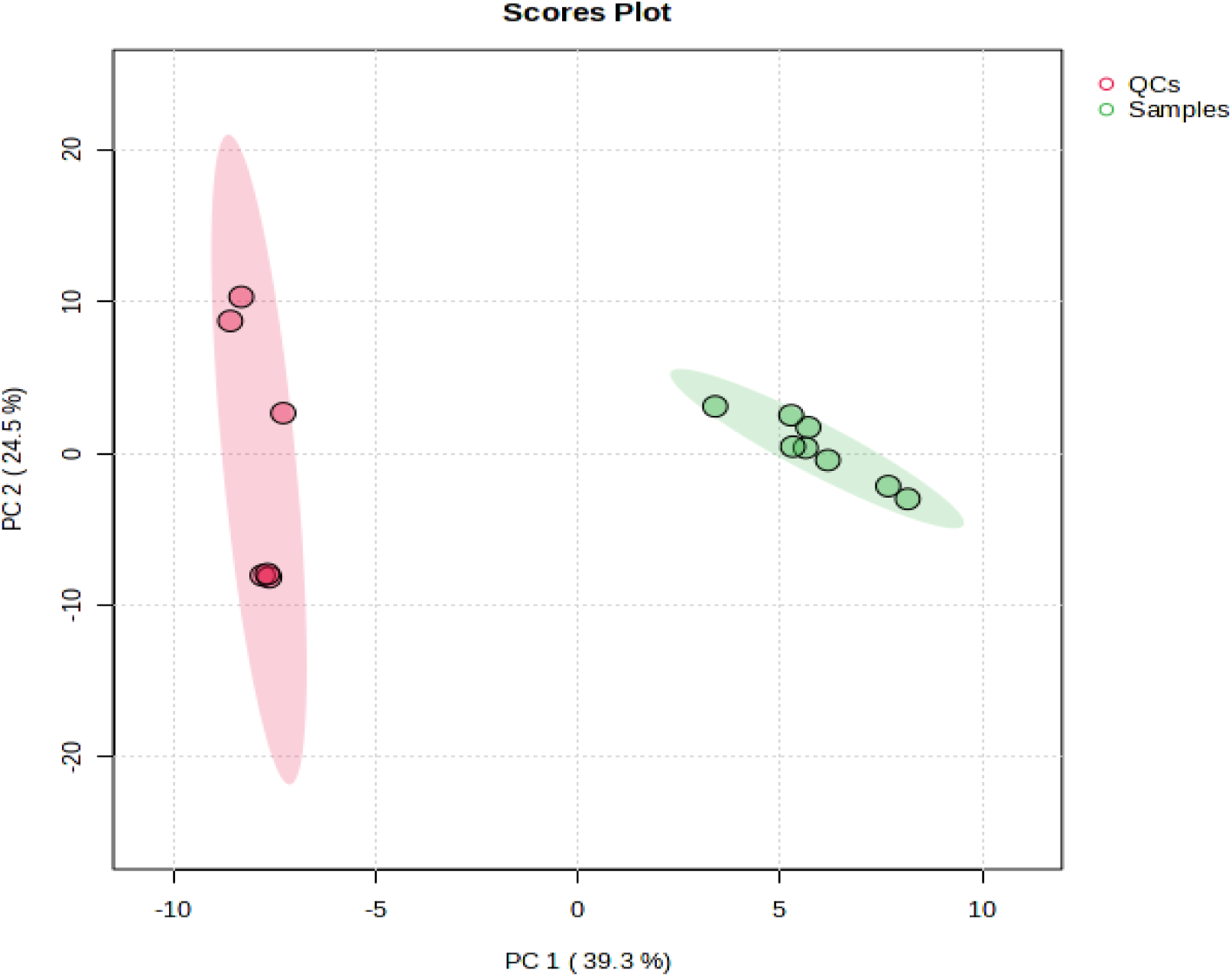
Principal component analysis scores plot showing distinct clustering of quality control samples (QCs) and separation between BALF experimental groups (Samples)

### Chemometric analysis result

Figure 2 shows an unbiased PCA model for all the metabolites included in CysLTR1 KO and WT samples groups. The x-axis (PC1) explains 33.3% of the total variance, while the y-axis (PC2) accounts for 23.7%.

**Figure 2.**
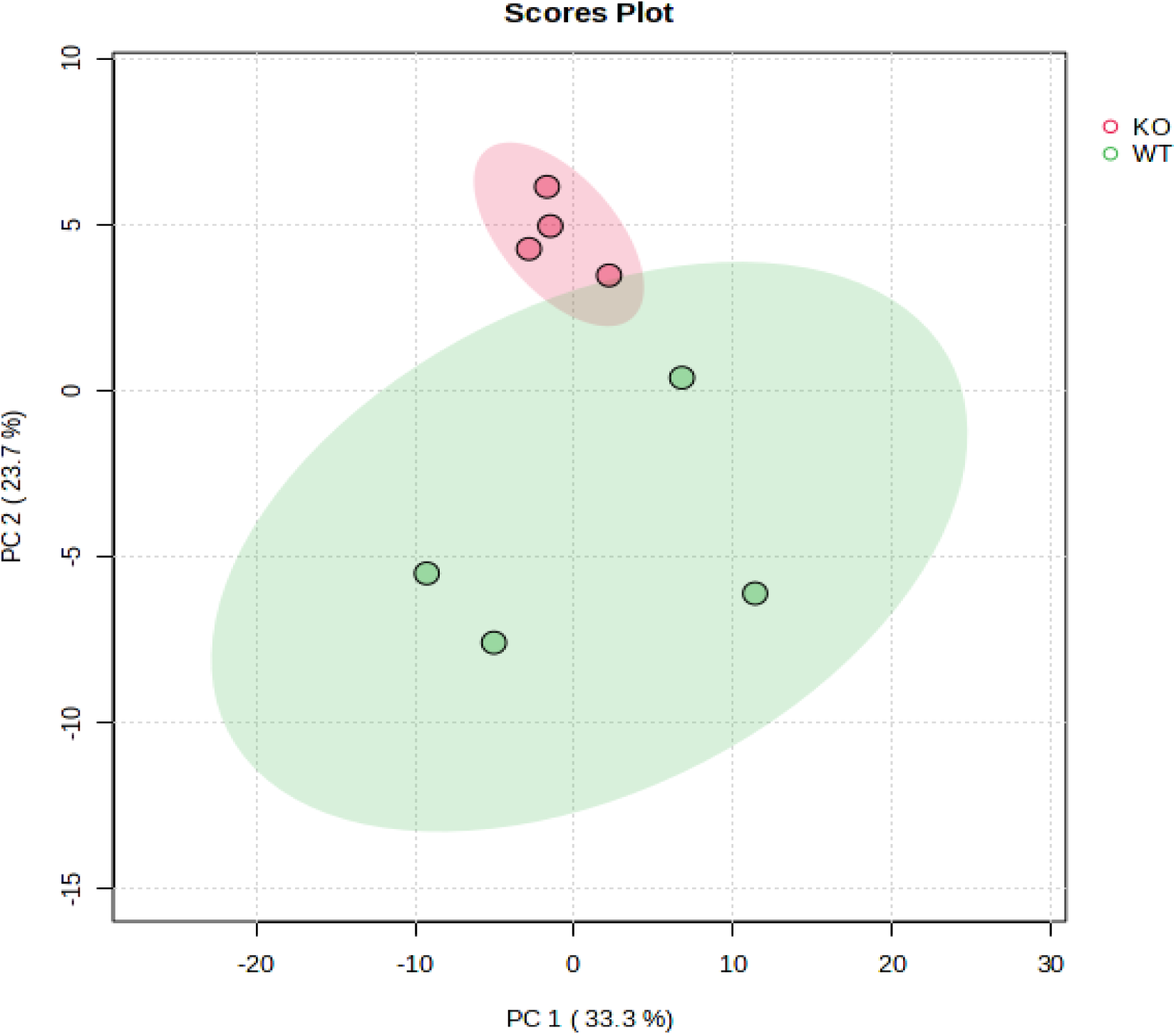
Principal component analysis scores plot for CysLT1 KO and WT samples groups. Percentages in brackets represent the proportion of variation explained in the observed data by the specific principal component (PC).

The clear separation between these groups along both PC1 and PC2 indicates significant metabolic differences between the compared CysLTR1 KO and WT groups. The ellipses further indicate that each group has distinct metabolic profiles with minimal overlap. This suggests that the absence of CysLTR1 has a strong effect on the overall metabolic variation of BALF samples when compared to that of the WT samples.

As shown in Figure 3, the OPLS-DA scores plot shows clear separation between CysLTR1 KO and WT groups. Each point represents an individual sample, with CysLTR1 KO samples (red circles) clustering on the left and WT samples (green triangles) clustering on the right. The x-axis (T score [1], 21.4%) captures the predictive variation between the groups, while the y-axis (orthogonal T score [1], 21.8%) represents variation within each group. The OPLS-DA variable of importance in projection (VIP) analysis identified 49 significant metabolites.

**Figure 3.**
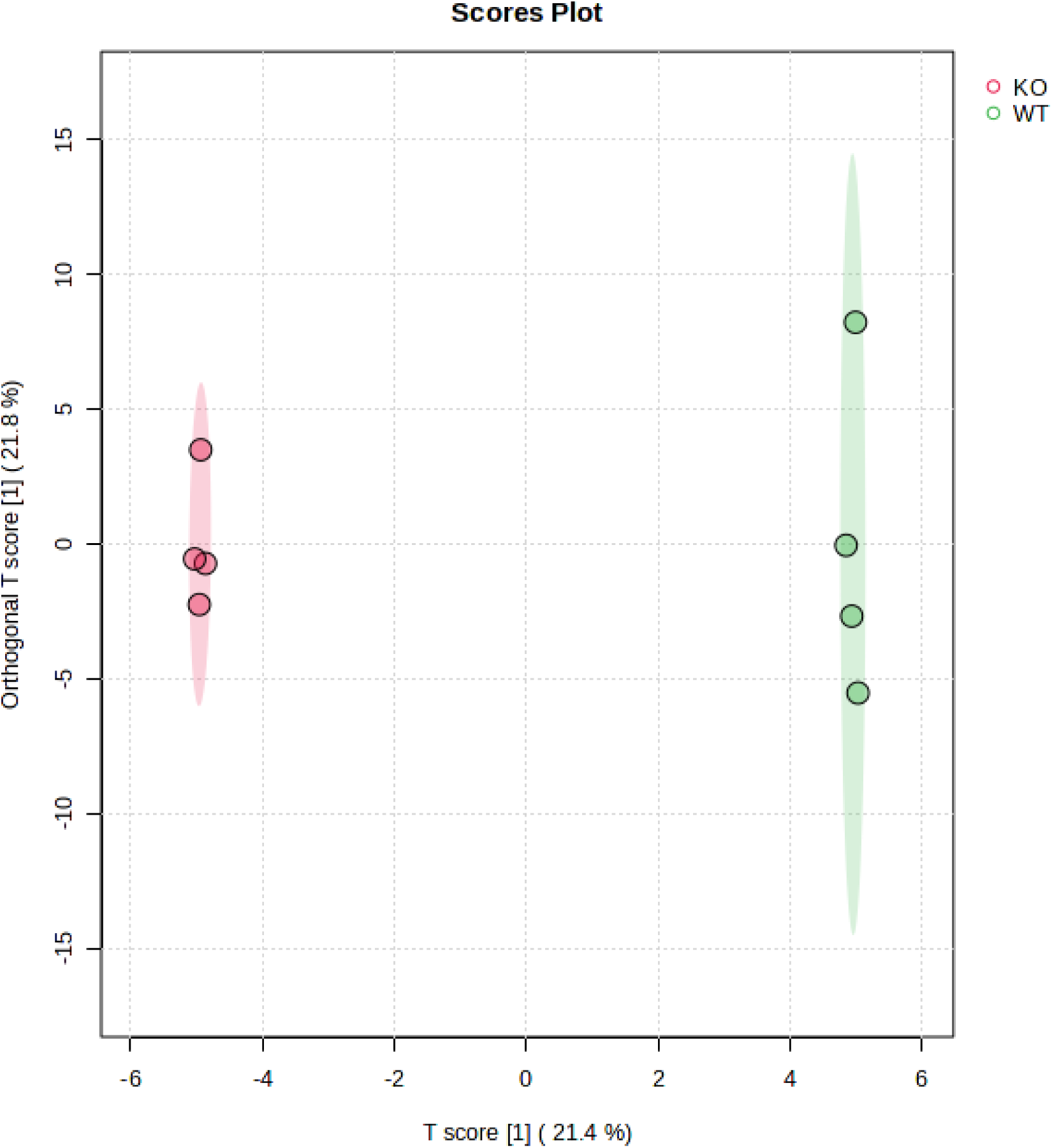
OPLS-DA scores plot showing distinct separation between CysLTR1 KO and WT BALF sample groups.

**Table 1.** OPLS-DA model performance using predictive and orthogonal components.

| Component | p1 (predictive) | o1 (orthogonal 1) | o2 (orthogonal 2) |
| --- | --- | --- | --- |
| **R<sup>2</sup>X** | 0.214 | 0.218 | 0.216 |
| **R<sup>2</sup>Y** | 0.922 | 0.0571 | 0.0204 |
| **Q<sup>2</sup>** | 0.576 | 0.154 | 0.0282 |

The table summarizes the performance of the OPLS-DA model, while explaining the variance and the predictive ability of its components. The predictive component (p1) shows a strong model fit, with an R²Y of 0.922, indicating that 92.2% of the variation in the response variable is explained by the model. The corresponding Q² value of 0.576 suggests acceptable predictive reliability, supporting the model’s ability to adapt to new data.

### Univariate and multivariate analysis result

Three univariate analyses were employed for variable selection: p-value t-test, fold change, and effect size. Applying a t-test with a p-value cut-ff of < 0.1 identified 23 significant metabolites. A log 2 fold change (Log2FC) analysis (threshold of 0.5), which assesses absolute differences between WT and KO group means, resulted in 70 metabolites. Effect size, with a threshold of Cohen’s d ≥ 0.8 was set, indicating a practically significant difference between groups identified 51 metabolites. The variable in projection (VIP) score from the orthogonal partial least-squares-discriminant analysis (OPLS-DA) was used as the multivariate analysis to identify significant metabolites, revealing a total of 49 metabolites.

### Marker selection

A total of 18 metabolite markers that best describe the variance between CysLTR1 KO deficient and WT BALF sample groups were selected using a multi-statistical selection approach based on the following criteria: a compound with t-test p-value < 0.1, a Log2FC threshold of 0.5, an effect size ≥ 0.8 and OPLSDA VIP > 1 was considered significant for further biological investigation.

Figure 4 below shows a Venn diagram for the selection of the metabolite markers showing most significance differences between the compared groups.

**Figure 4.**
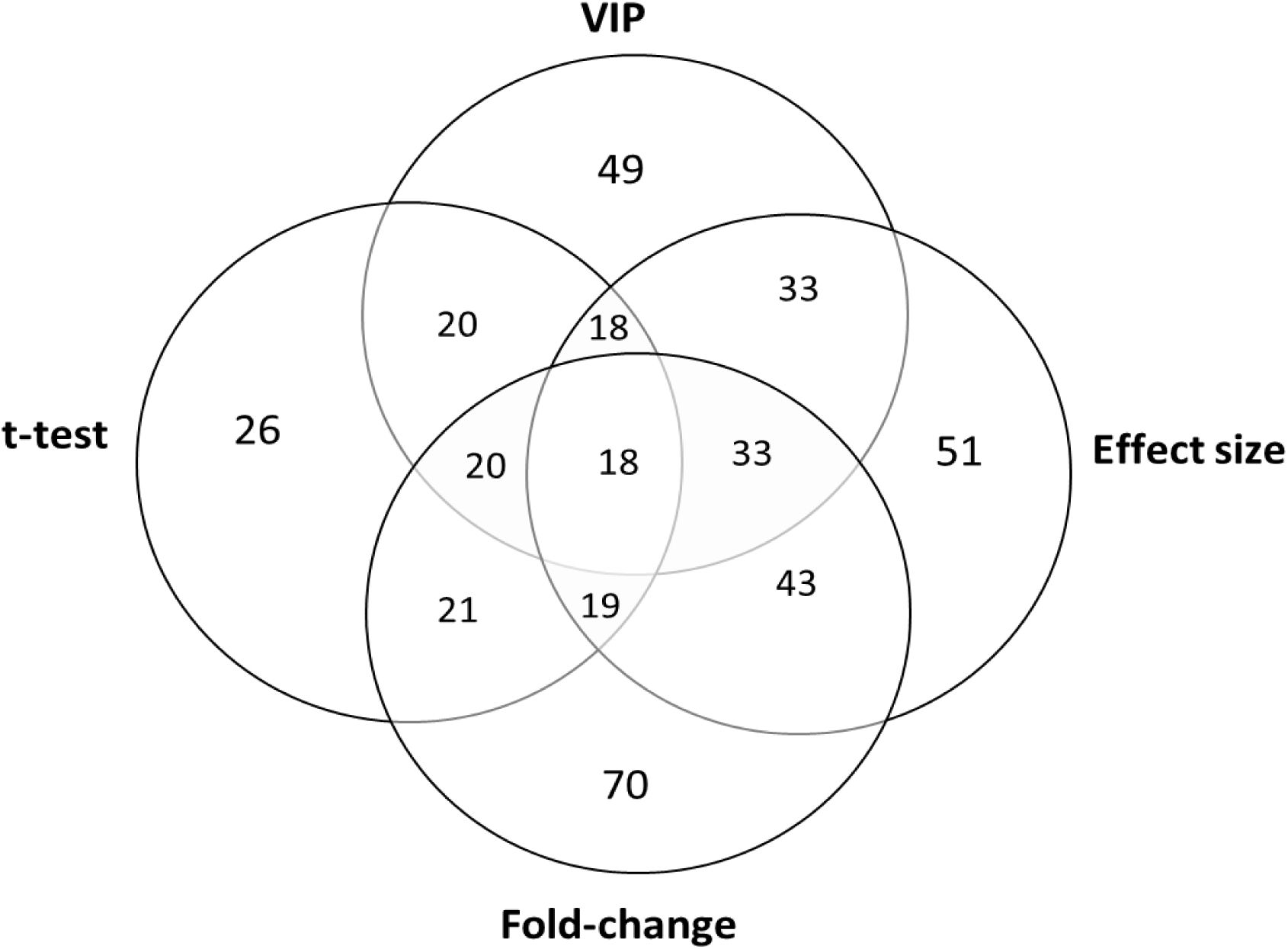
Venn diagram indicating compound selection of the 18 metabolite markers contributing most significantly to the variation between the CysLTR1-KO and WT BALF sample groups using a multi-statistical approach. Selection criteria included: having a t-test p-value < 0.1, a Log2FC 0.5, OPLSDA VIP > 1 and an effect size ≥ 0.8.

**Table 2.** Metabolites best describing the variance between the CysLTR1 KO and WT BALF mice sample groups.

| Metabolite<br>(PubChem ID) | Average concentration (ng/L)<br>(standard deviation) |  | Log2FC<br>(threshold<br>0.5) | Cohens<br>(effect size:<br>d-value ≥ 0.8) | Vip > 1 | t-test<br>p-value<br>< 0.1 |
| --- | --- | --- | --- | --- | --- | --- |
|  | Wild type mice | CysLTR1 KO mice |  |  |  |  |
| Tricarballic acid<br>(14925) | 2.650<br>(1.994) | 0.792<br>(0.100)↓ | -4.0294 | 2.911 | 1.993 | 0.001 |
| Glucosamine (739) | 9,707<br>(10.121) | 5,825<br>(8.381)↓ | -2.45 | 1.958 | 1.658 | 0.026 |
| Decan-2-ol (2737541) | 1.117<br>(0.987) | 0.120<br>(0.269)↓ | -2.4346 | 1.945 | 1.647 | 0.027 |
| D-Chiro-Inositol<br>(16216949) | 125.524<br>(60.474) | 56.153<br>(14.225)↓ | -1.639 | 2.558 | 1.987 | 0.013 |
| Glucose (5793) | 2.118<br>(1.647) | 0.780<br>(0.593)↓ | -1.4403 | 0.824 | 1.793 | 0.036 |
| Mandelic acid<br>(22943025) | 52.300<br>(18.357) | 38.746<br>(37.000)↓ | -1.3661 | 3.362 | 1.946 | 0.002 |
| Dodecane (8182) | 6.909<br>(5.539) | 3.410<br>(3.176)↓ | -1.6753 | 1.493 | 1.393 | 0.082 |
| D-xylose (135191) | 23.682<br>(19.270) | 9.937<br>0.00212097↓ | -1.7129 | 1.307 | 1.619 | 0.033 |
| Glutaric acid<br>(90037731) | 37.290<br>(17.780) | 20.310<br>5.905↓ | -1.3434 | 2.186 | 1.828 | 0.008 |
| Sedoheptulose<br>(5459879) | 19.677<br>(6.298) | 12.138<br>(4.172)↓ | -1.1911 | 2.803 | 1.967 | 0.002 |
| Glyceric acid (752) | 55.715<br>(12.841) | 31.324<br>(7.776)↓ | -1.1739 | 2.939 | 1.955 | 0.002 |
| Gluconic acid (10690) | 222.814<br>(134.281) | 155.146<br>(80.397)↓ | -1.4347 | 1.624 | 1.690 | 0.022 |
| 1-Hexadecanol (2682) | 38.672<br>(19.981) | 22.849<br>(9.901)↓ | -1.2736 | 1.758 | 1.797 | 0.011 |
| (R*,S*)-2,3 Dihydroxy<br>butanoic acid (250402) | 8.782<br>(3.632) | 5.921<br>(1.384)↓ | -1.1014 | 2.605 | 1.848 | 0.006 |
| 1-Methylinosine<br>(65095) | 0.837<br>(0.572) | 0.555<br>(0.265)↓ | -1.1817 | 1.643 | 1.394 | 0.088 |
| Butanoic acid<br>(16213394) | 325.566<br>(117.085) | 207.031<br>(36.553)↓ | -0.98067 | 2.737 | 1.836 | 0.007 |
| Myo-inositol (440388) | 1912.119 | 1398.837 | -0.93205 | 1.676 | 1.593 | 0.037 |
|  | (865.363) | (441.522)↓ |  |  |  |  |
| Galactose (441035) | 25632.153<br>(6700.214) | 20646.841<br>(2718.550)↓ | -0.63269 | 2.766 | 1.907 | 0.003 |

## Discussion

Figure 5 below shows an overview of the interconnected metabolic pathways showing metabolites (highlighted in green) detected to be significantly altered in the CysLTR1 knockout mice BALF samples when compared to the WT mice. Altered pathways in the CysLTR1 knockout mice include changes to glycolysis, the pentose phosphate pathway, purine metabolism, amino acid metabolism, fatty acid metabolism, galactose metabolism, and the hexosamine biosynthesis pathway. The schematic diagram illustrates the metabolic concentration changes associated with CysLTR1 deficiency.

**Figure 5.**
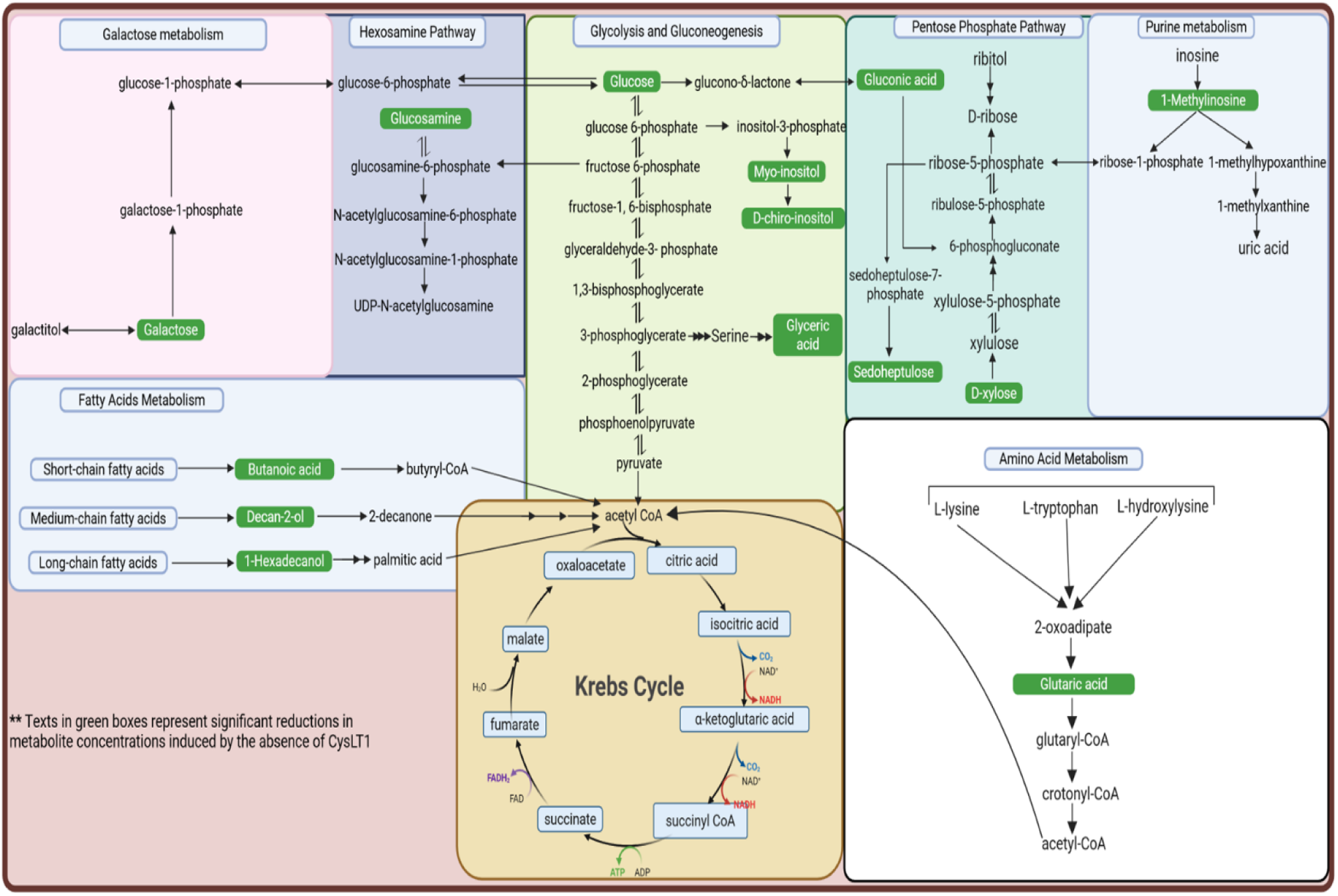
An overview of the integrated metabolic pathways of altered metabolites (arrows show directional changes in concentrations) in BALF samples of CysLTR1 KO mice when compared to that of the WT mice.

The metabolomic analysis of biofluids such as serum, plasma, urine, cerebrospinal fluid, and BALF, is a powerful tool for understanding the physiological and pathological states of disease conditions, and in this case lung health (13,14). In clinical research, information providing a comprehensive snapshot of the metabolic alterations in a biological system, is considered important for identifying disease biomarkers, elucidating altered biochemical pathways, and monitoring therapeutic responses (15,16). In the present investigation, the results provide useful insights into the biochemical and physiological perturbations associated with a CysLT1 deficiency, further highlighting the potential role of CysLT1 in pulmonary physiology and disease mechanisms, as revealed by changes in the metabolic pathways influenced by the absence of CysLTR1 in the KO mice (Figure 5).

The most important finding of this study is the significant reduction in glucose levels in the BALF of CysLT1 KO mice, confirming a previously reported association between cysteinyl leukotriene signalling and pulmonary glucose homeostasis (3,4). Furthermore, (17) reported the negative regulation of glucose-stimulated insulin secretion in MIN6 cells; a mouse pancreatic beta-cell line by CysLTR1 receptors. In the absence of CysLTR1, insulin secretion is improved, and hence an improved systemic glucose clearance and glucose uptake into insulin dependent tissues including skeletal muscle, adipose tissue and the heart. This would result in a reduced glucose availability to other organs that take up glucose via non-insulin dependent pathways, including the lung and hence result in the reduced glucose levels in the BALF of the CysLTR1 KO mice. This observation provides important insight into the role of CysLTR1 in regulating the metabolic microenvironment of the lung, particularly under inflammatory conditions. CysLTR1, a receptor for leukotrienes LTC₄, LTD₄, LTE₄ and LTB₄ (18), plays a critical role in mediating inflammatory responses in the lung (19,20). The absence of CysLTR1 disrupts the leukotriene-mediated signalling cascade, resulting in reduced inflammatory cell response in the airways. Since the activation of immune cells such as M1 macrophages, neutrophils and T lymphocytes causes metabolic reprogramming, and a heavy reliance on glycolysis which subsequently consumes substantial amounts of glucose during inflammation (21,22), the observed reduction in glucose levels in the BALF of CysLTR1 KO mice may therefore also be attributed to the decline in basal pulmonary inflammatory response resulting from the disruption of leukotriene-mediated signalling. Given that CysLTR1 is essential for activating inflammatory cells in the airways through the lipid-mediated actions of leukotrienes, its absence therefore likely reduces the presence and activity of these cells, leading to decreased pulmonary glucose uptake and consumption.

Although glucose is a fundamental energy source and a key metabolite in glycolysis, it also serves as a substrate for the pentose phosphate pathway (PPP), hexosamine pathways, and the tricarboxylic acid (TCA) cycle. Hence, the reduced levels of glucose in this study, in turn leads to downstream metabolic perturbations, as seen in the downregulation of several carbohydrate-related metabolite intermediates including myo-inositol, D-chiro-inositol, glyceric acid, glucosamine, gluconic acid, and sedoheptulose, all of which are directly or indirectly linked to glucose metabolism.

Myo-inositol and D-chiro-inositol are isomeric forms of inositol, a sugar alcohol that plays a role in cellular growth and insulin processing (23). Both metabolites are synthesized from glucose-6-phosphate as seen in figure 5. In the absence of adequate glucose supply, the availability of glucose-6-phosphate is limited, leading to reduced inositol biosynthesis (24). Since myo-inositol is the precursor of D-chiro-inositol, a further reduction in the latter supports this finding. The depletion of these inositols not only indicates an impaired carbohydrate metabolism (24), but also insulin signal transduction (23) and membrane phospholipid composition (25), particularly in tissues such as the lung and liver. Their role in regulating pathways related to type 2 diabetes mellitus has been widely investigated. For instance, myo-inositol has been reported to inhibit glucose absorption in the intestine and promote muscle glucose uptake in rats, while clinical trials have demonstrated that both myo-inositol and D-chiro-inositol possess insulin-mimetic properties and an ability to improve insulin sensitivity in metabolic conditions associated with insulin resistance in humans (26,27). As mentioned, glyceric acid is also a downstream metabolite of glycolysis (Figure 5). Reduced glucose supply impairs glycolytic throughput, which ultimately results in the reduced production of its glycolytic intermediates, 3-phosphoglycerate and glyceraldehyde-3-phosphate. Consequently, the downstream production of glyceric acid is diminished. The reduced levels of glyceric acid, further supports disruption in the glycolytic pathway, with an implication on glucose utilization and the metabolic adaptability in the CysLTR1 KO mice.

Glucosamine was also reduced in the BALF of the CysLT1 KO group comparatively, most likely due to a reduction in the availability of its precursor glucose. Glucosamine in turn is a precursor for glycosaminoglycans and glycoprotein synthesis, macromolecules that play crucial roles in various biological processes, including cell signalling, tissue development, and disease progression (28,29,30,31). It’s downregulation in the CysLTR1 KO group can result in an impairment in protein glycosylation. This may impact pulmonary function, as glycosylation is crucial for maintaining mucosal barrier integrity and immune responses. Since the monosaccharides glucose and galactose share a close metabolic relationship, it is expected that a reduction in the levels of glucose likely contributes to a corresponding decrease in galactose production.

Although the production of gluconic acid, from the oxidation of glucose, is not a primary metabolic route for glucose metabolism in mammals, key enzymes of the shunt such as glucose dehydrogenase and gluconate kinase (32) have been identified in mammals. Additionally, gluconic acid can be produced via the gluconate shunt pathway, through gut microbial activity (33). Due to involvement of CysLTR1 in modulating inflammatory signalling and its putative role in indirectly influencing glucose metabolism, including pathways such as glycolysis, PPP, and oxidative phosphorylation, the reduced levels of gluconic acid in the BALF of CysLTR1 KO mice may be associated with decreased glucose availability. This condition can downregulate glucose oxidase activity, thereby contributing to lower gluconic acid levels (34).

The PPP plays a critical role in generating nucleotide precursors such as ribose-5-phosphate (Figure 5), and for maintaining a dynamic balance between NADP⁺ and NADPH synthesis (35,36). CysLTR1 in BALF of KO mice leads to impaired cysteinyl leukotriene signalling, which ultimately results in reduced glucose levels in the pulmonary environment as previously explained. Since inflammatory cells rely on the PPP to support rapid proliferation and anabolic metabolism, reduced inflammation and glucose supply to PPP will result in diminished PPP flux, consistent with the reduction in some PPP-associated metabolites such as D-xylose, gluconic acid and sedoheptulose. D-xylose is metabolized through the non-oxidative branch of the PPP, where it is interconverted via intermediates such as xylulose-5-phosphate to eventually produce ribose-5-phosphate. Similarly, gluconic acid (derived from glucose) also serves as a precursor for ribose-5-phosphate (35,36). The reduced availability of glucose, D-xylose and their resulting PPP precursors limits the capacity of the PPP to generate ribose-5-phosphate, required for nucleotide synthesis. This impairment in PPP metabolic activity is further supported in our study by the reduction in sedoheptulose in the CysLTR1 KO mice. Sedoheptulose, an intermediate in the PPP, plays a crucial role in metabolism by contributing to NADPH generation primarily for reductive biosynthesis, and to ribose formation for nucleotide biosynthesis (37). This resulting decrease in nucleotide synthesis is further substantiated by the reduced levels of 1-methylinosine, a methylated purine nucleoside involved in purine metabolism in the CysLTR1 KO mice of our study. This inevitably results in RNA metabolism or turnover dysregulation, which would have downstream effects on gene expression and protein synthesis.

1-Hexadecanol and butanoic acid, both byproducts of fatty acid metabolism (Figure 5) are also reduced in the CysLTR1 KO mice. This may reflect a metabolic adaptation to compensate for lower glucose availability, thereby increasing utilization of fatty acid derivatives for energy in the lung. Furthermore, a reduction in dodecane levels reflect altered lipid metabolism, resulting from disrupted leukotriene signalling in the pulmonary environment associated with CysLTR1 deficiency. Leukotrienes, which are lipid mediators derived from arachidonic acid (38), play a key role in inflammatory processes, and disruptions in arachidonic acid pathways have been associated to changes in lipid metabolism (39). Given the past report of microbial biosynthesis of alkanes from fatty acids (40), another plausible explanation is that CysLTR1 deficiency may alter the pulmonary microbiota, thereby reducing microbial conversion of endogenous fatty acids into dodecane.

Glutaric acid is a metabolic product of lysine and tryptophan metabolism (42). These amino acids are broken down into glutaryl-CoA, which is converted into crotonyl-CoA and eventually enters the TCA cycle (43). The reduction in mandelic acid and the aforementioned glutaric acid are both also indicative of an altered intestinal microbiome (44,45). Recent studies have linked both metabolites to microbial activity and dysbiosis in inflammatory conditions (44,46). Thus, the reduced levels of these metabolites in CysLTR1 KO mice reflects an interplay between the host metabolic deregulation, inflammatory response and microbial metabolic shifts. Furthermore, we also observed a notable reduction in the concentrations of tricarballylic acid (TA) and decan-2-ol in the BALF samples of CysLTR1 KO mice when compared to the WT controls. TA and decan-2-ol are associated with corn-derived feed (47). TA is known to be a microbial fermentation product formed by rumen microorganisms through the conversion of trans-aconitic acid (TAA), a compound abundant in corn and accounting for approximately 95% of its total aconitic acid (48). Although no direct link between CysLTR1 and the gut or intestinal microbiota has been reported in particular, G protein-coupled receptors, the most abundant class of cell surface receptors in the human body and to which CysLTR1 belongs, are known to interact with the gut microbiota, influencing various physiological processes (49). Therefore, the observed reduction in these metabolites most likely result from changes in microbial metabolism associated with the loss of CysLTR1-mediated inflammatory signalling, which may indirectly modulate the host–microbiome interface. Furthermore, The deletion of CysLTR1 could lead to diminished immune response, thereby altering the microbial niche and suggesting that microbial communities capable of this transformation may either be reduced in number or functionally inactive. Aside the host metabolism, the immunological shifts which resulted in the observed metabolic changes in the BALF of CysLTR1 KO mice may also alter the respiratory microbiome. Inflammatory mediators including leukotrienes have been reported to influence the composition and activity of microbial communities (50).

## Conclusion

In this study, we utilized GC-MS-based metabolomics approach to investigate the metabolic alterations in the bronchoalveolar lavage fluid (BALF) of CysLTR1 knockout (KO) mice. The study provides valuable insights into the biochemical and physiological consequences of CysLTR1 deficiency in the mammalian pulmonary environment and the role of CysLTR1 in the lung. Many key metabolic pathways necessary for maintaining pulmonary homeostasis were affected, including the metabolism of carbohydrates, lipids, amino acids, nucleotides and the lung microbiome, providing compelling evidence that the deletion of CysLTR1 significantly alters pulmonary metabolic homeostasis.

The observed changes in carbohydrate, lipid, nucleotide, and amino acid metabolism underscore the central role of CysLT1 in insulin signalling and inflammatory response as suggested by previous studies (17), as well as maintaining metabolic balance in lung tissue. Additionally, the downregulation of PPP intermediates in CysLTR1 KO indicates impaired production of NADPH and ribose-5-phosphate, which are essential for supporting nucleotide synthesis (35).

Generally, these findings highlight the use of BALF metabolomics as a sensitive approach to determine localized metabolic alterations and identify molecular signatures associated with CysLTR1 deletion in the lung. This study provides a foundation for future mechanistic studies into CysLTR1 as a potential therapeutic target for metabolic and respiratory disorders.

## Notes

### Competing Interest Statement

The authors have declared no competing interest.

### Funding

- National Research Foundation (NRF) of South Africa: SRUG2204062208, RDYR180413320675, IFR180305315866, RA22110567937, SRUG22051611051
- South African Medical Research Council (SAMRC)
- Fogarty International Centre of the National Institutes of Health under Award Number: K43TW012587

